# Human aged astrocytes induce neurotoxicity in response to inflammatory stimuli

**DOI:** 10.1101/2025.08.17.670762

**Authors:** Hayato Kobayashi, Kiyohiro Maeda, Takashi Wakui, Kenichi Kazetani, Akira Nabetani, Euikyung Shin, Hiroshi Kato, Setsu Endoh-Yamagami

## Abstract

Astrocytes play a critical role in neuroinflammation and the pathogenesis of neurodegenerative diseases. Here we found that human induced pluripotent stem cell (iPSC)-derived astrocytes responded differently to inflammatory triggers compared to rodent astrocytes, showing increased neurotoxicity when exposed to TNF-α and IFN-γ. Furthermore, astrocytes with senescent features showed even higher levels of neurotoxicity in the presence of TNF-α and IFN-γ, suggesting a potential link between aging and neurodegenerative diseases. It was also demonstrated that LPS-activated neuron/astrocyte/microglia tri-culture produced TNF-α, leading to neurotoxicity in the tri-culture when IFN-γ was present. Through compound screening, we identified Janus kinase inhibitors capable of preventing neurotoxicity in astrocytes induced by TNF-α and IFN-γ, demonstrating the potential use of neurotoxic astrocytes as a platform for drug screening. These results provide insight into the complex relationship between aging, inflammation, and neurodegenerative diseases, emphasizing the potential of targeting astrocytes as a novel therapeutic approach for addressing neurodegenerative diseases.

## Main

Astrocytes play a central role in neuroinflammation and the etiology of neurodegenerative diseases, in addition to their well-known supportive functions to neurons ^1,2^. Liddelow and Barres’s group first reported that neurotoxic reactive astrocytes are induced by the signaling molecules IL-1α, tumor necrosis factor (TNF), and complement component 1q (C1q) secreted by activated microglia using rodent primary cells ^1^. The neurotoxic reactive astrocytes exhibit hypertrophic morphology and express high levels of complement component 3 (C3) and other neurotoxic reactive astrocyte markers (e.g. *GBP2*, *PSMB8*, *FBLN5*, *SERPING1*) along with pan-reactive astrocyte markers (e.g. *LCN2*, *STEAP4*, *CXCL10*) ^1^. An increased number of C3-positive astrocytes have been observed in disease model mice and human patients with neurodegenerative diseases, including Alzheimer’s disease (AD), Parkinson’s disease (PD), and amyotrophic lateral sclerosis (ALS) ^1,3–5^. It has been reported that astrocytes prepared from postmortem tissues of ALS patients show neurotoxicity ^6^, consistent with findings indicating that primary astrocytes isolated from aged and/or ALS mice have reduced support for neurons ^7^. Although the involvement of neurotoxic reactive astrocytes in neurodegenerative diseases has been implicated in both rodents and humans, the majority of in vitro studies on neurotoxic reactive astrocytes utilize rodent primary astrocytes, likely due to the difficulty of obtaining primary cells from human tissues.

Here we utilized human induced pluripotent stem cell (iPSC)-derived astrocytes and demonstrated that human astrocytes exhibited pronounced neurotoxicity in response to TNF-α and IFN-γ, different factors reported in rodent studies. In addition, astrocytes with senescent features showed increased neurotoxicity in response to TNF-α and IFN-γ, prompting speculation about relationship between aging and the development of neurodegenerative diseases. Furthermore, the identification of compounds with activities to inhibit neurotoxicity through screening suggests the potential usage of the screening system for drug development.

## Results

### Combination of TNF-α and IFN-γ induced neurotoxicity in human iPSC-derived astrocytes

To induce neurotoxicity in human iPSC-derived astrocytes, we first treated the astrocytes with IL-1α, TNF-α, and C1q (Fig. 1a). However, these three factors did not induce the characteristic morphological changes from star-shaped to hypertrophic morphology as reported in rodent astrocytes ^1^. We hypothesized that this result could be attributed to differences in cytokine response between rodent and human astrocytes. Then, we focused on IL-1β and IFN-γ, which are known to activate human astrocytes but not mouse astrocytes ^8^. By adding IL-1β and IFN-γ to the three factors, astrocytes showed morphological changes (Fig. 1a). Upon removing each factor individually, we found that the combination of TNF-α and IFN-γ was sufficient to induce these morphological changes (Fig. 1a, b).

**Fig. 1.**
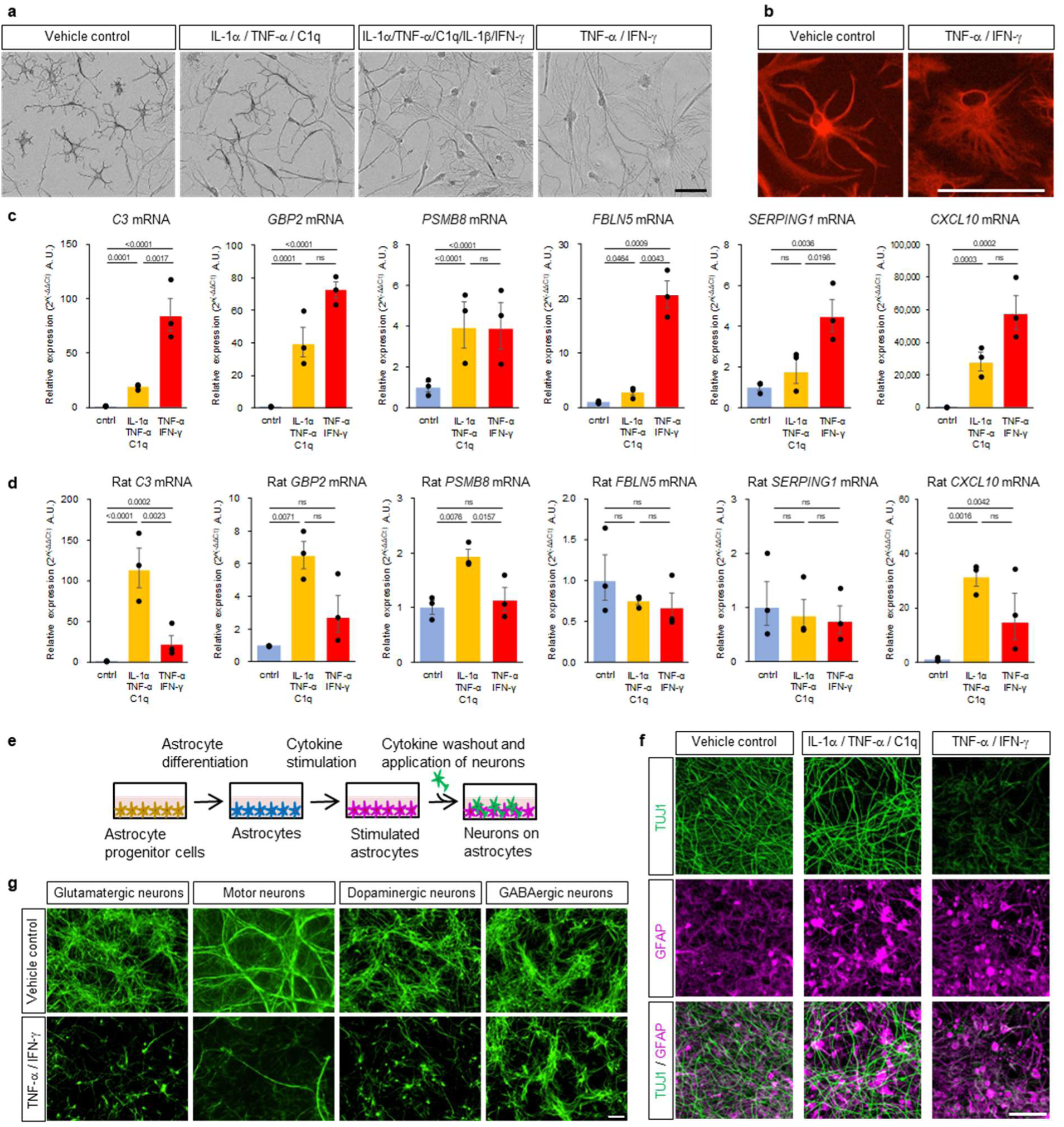
| Combination of TNF-α and IFN-γ induces a reactive neurotoxic state in human iPSC-derived astrocytes. **a,** Morphology of human astrocytes treated with IL-lα/TNF-α/Clq (3.0. 30. 400 ng/mL). IL-lα/TNF-α/Clq/IL-lβ/ IFN-γ (3.0.30. 400. 2.5,5.0 ng/mL), or TNF-α/IFN-γ (7.5. 5.0 ng/mL) for two days, **b,** Immunostaining of human astrocytes heated with TNF-α/IFN-γ using GFAP antibodies, **c-d,** Relative gene expression levels of *C3, GBP2, PSMBS, FBLN5, SERPING1.* and *CXCL10* using *GAPDH* as the internal control hi human iPSC-derived astrocytes **(c)** and in rat primary astrocytes **(d)** cultured in differentiation medium. Bar data represent mean ± s.e.m.; each dot represents an individual biological replicate (π = 3 independently prepared cells). Statistical analysis was performed using two-way ANOVA followed by Tukey’s multiple comparisons test; ns: 0.05 < *P.* **e,** Schematic of neurotoxicity assay flow, **f**, Evaluation of neurotoxicity of astrocytes treated with IL-lα/TNF-α/Clq or TNF-α/IFN-γ using Ngn2-induced cortical excitatory neurons, **g,** Neurotoxicity against glutamatergic, motor, dopaminergic, and GABAergic neurons. Scale bar: 100 µm **(a, b, f, g)**

The combination of TNF-α and IFN-γ also induced the neurotoxic astrocyte marker genes (*C3*, *GBP2*, *PSMB8*, *FBLN5*, *SERPING1*) and pan-reactive astrocyte marker *CXCL10* more strongly in human iPSC-derived astrocytes compared to the combination of IL-1α, TNF-α, and C1q (Fig. 1c). In contrast, the combination of the three factors (IL-1α, TNF-α, and C1q) induced higher expression of these neurotoxic and pan-reactive marker genes in rat primary astrocytes compared to the combination of TNF-α and IFN-γ (Fig. 1d). This pattern was consistent across different culture media (Extended Data Fig. 1a, b), supporting species-specific differences in astrocyte cytokine responses. Elevated levels of C3 were also observed in exosomes isolated from the culture medium of astrocytes treated with TNF-α and IFN-γ (Extended Data Fig. 1c), which is consistent with the increased levels of C3 in astrocyte-derived exosomes in AD patients ^9^.

To examine the neurotoxicity of cytokine-treated astrocytes, NGN2-induced cortical excitatory neurons were seeded on astrocytes following a medium washout (Fig. 1e, f). Neurite staining with TUJ1 antibodies revealed more severe neuronal damage induced by the combination of TNF-α and IFN-γ compared to the three-factor combination. The impact of the neurotoxicity was compared by culturing different types of neurons on astrocytes, revealing that excitatory neurons, including glutamatergic neurons, motor neurons, and dopaminergic neurons, were more sensitive to neurotoxic astrocytes compared to inhibitory GABAergic neurons (Fig. 1g).

### Compound screen identified JAK inhibitors that suppressed the acquisition of neurotoxicity in astrocytes

To explore the potential use of our neurotoxic human astrocyte platform for therapeutic development, we conducted a screen to identify compounds that suppress the acquisition of neurotoxicity. Human iPSC-derived astrocytes were treated with 5,301 bioactive small molecules in combination with TNF-α and IFN-γ for 24 hours, followed by RNA extraction and quantitative analysis of *C3* gene expression. The compounds that inhibited C3 expression 95% or more compared to the vehicle control at 10 μmol/L were selected as primary hits (Fig. 2a).

**Fig. 2.**
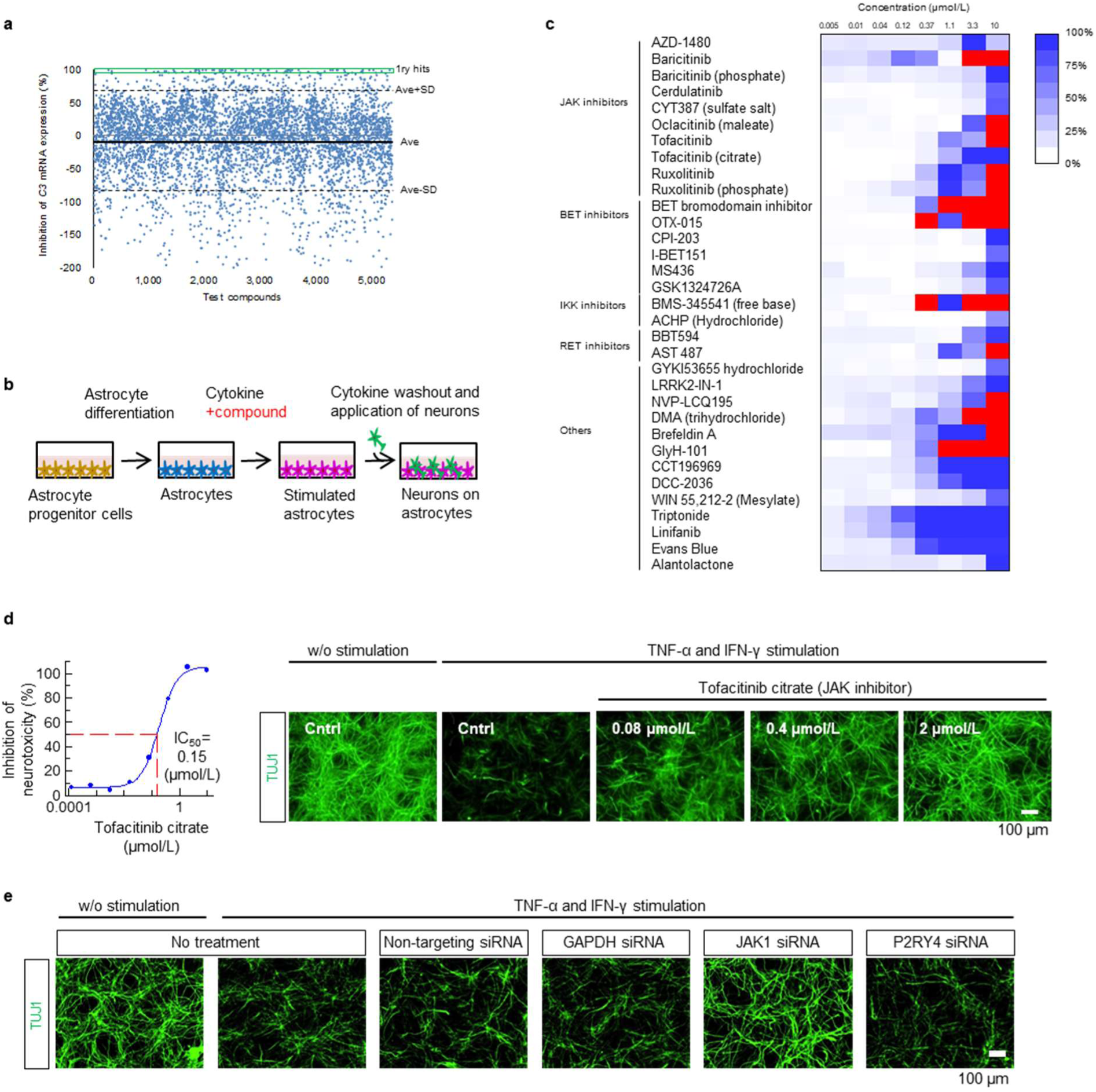
| Phenotypic screening reveals the involvement of JAK in the acquisition of neurotoxicity. **a,** Scatter plot of the primary screen evaluated for the activity of repressing *C3* mRNA expression, **b,** Schematic of the neurotoxicity assay for compounds, **c,** Dose-response of hit compounds with activity to inhibit the acquisition of neurotoxicity evaluated in NGN2-induced cortical excitatory neurons. The data are presented as the mean of *n*=2 technical replicates. The reproducibility of the data was confirmed in an independent experiment, **d,** Representative data of neurotoxicity assay of astrocytes treated with Tofacitinib citrate using NGN2-induced cortical excitatory neurons. The dose-responsive curve is presented as the mean of *n =* 2 technical replicates, **e,** Neurotoxicity assay of astrocytes treated with siRNA against *GAPDH, JAKI,* or *P2RY4.* Scale bar: 100 µm **(d, e)**

To further evaluate the ability of compounds to inhibit the acquisition of neurotoxicity, we developed a method to quantify neurite length. This involved immunostaining neuron-astrocyte co-cultures with TUJ1 antibodies, followed by measuring TUJ1-positive neurite length. The method successfully quantified the activity of blocking antibodies against TNF-α and IFN-γ in preventing the acquisition of neurotoxicity in astrocytes (Extended Data Fig. 2a). The primary hit compounds were applied to astrocyte cultures along with TNF-α and IFN-γ for 48 hours. The culture media containing the compounds and cytokines were then washed out, and NGN2-induced cortical excitatory neurons were seeded on the astrocytes, followed by a 72-hour incubation. Neurite length in the co-culture was subsequently quantified to identify and validate compounds with activity in inhibiting the acquisition of neurotoxicity (Fig. 2b, c). Multiple hits were identified among the Janus kinase (JAK) inhibitors (10 out of 32 validated hits), followed by BET inhibitors (6/32), IKK inhibitors (2/32), and RET inhibitors (2/32).

We further confirmed that the effects of the JAK inhibitor on astrocytes were effective across various neuronal types, including dopaminergic neurons and motor neurons in addition to excitatory cortical neurons (Fig. 2d and Extended Data Fig. 2b). To validate the target of JAK inhibitors, we performed siRNA knockdown against JAK kinases (Fig. 2e and Extended Data Fig. 2c). JAK1-specific siRNA suppressed the acquisition of neurotoxicity in astrocytes, whereas non-targeting siRNA, GAPDH siRNA, and P2RY4 siRNA had no effect. The knockdown of *JAK2* and *JAK3* resulted in the detachment of astrocytes from the culture plate (data not shown), which prevented the evaluation of their involvement in the acquisition of neurotoxicity. These results suggested that inhibition of JAK1 alone was sufficient to inhibit the acquisition of neurotoxicity and support the notion that the effects of JAK inhibitors are mediated through their supposed target.

### LPS-activated microglia induced neurotoxicity of astrocytes in the presence of IFN-γ

Microglia play a central role in neuroinflammation, and they are known to produce inflammatory cytokines in response to stimuli, including lipopolysaccharides (LPS) ^10^. To examine whether LPS-activated microglia are sufficient to induce neurotoxicity in astrocytes, we treated neuron/astrocyte co-cultures and neuron/astrocyte/microglia tri-cultures with LPS (Fig. 3a, b). LPS treatment alone did not induce neurotoxicity in either condition. We observed that the production of TNF-α was induced by LPS in human iPSC-derived neuron/astrocyte/microglia tri-culture (Fig. 3c) but the expression of IFN-γ was not detected (data not shown). Then, we treated neuron/astrocyte/microglia tri-cultures with a combination of LPS and IFN-γ and found that neurotoxicity was induced. In contrast, the combination of LPS and IFN-γ did not induce neurotoxicity in neuron/astrocyte co-cultures. These results suggest that TNF-α, released by LPS-activated microglia, induced astrocyte-mediated neurotoxicity in the presence of IFN-γ.

**Fig. 3.**
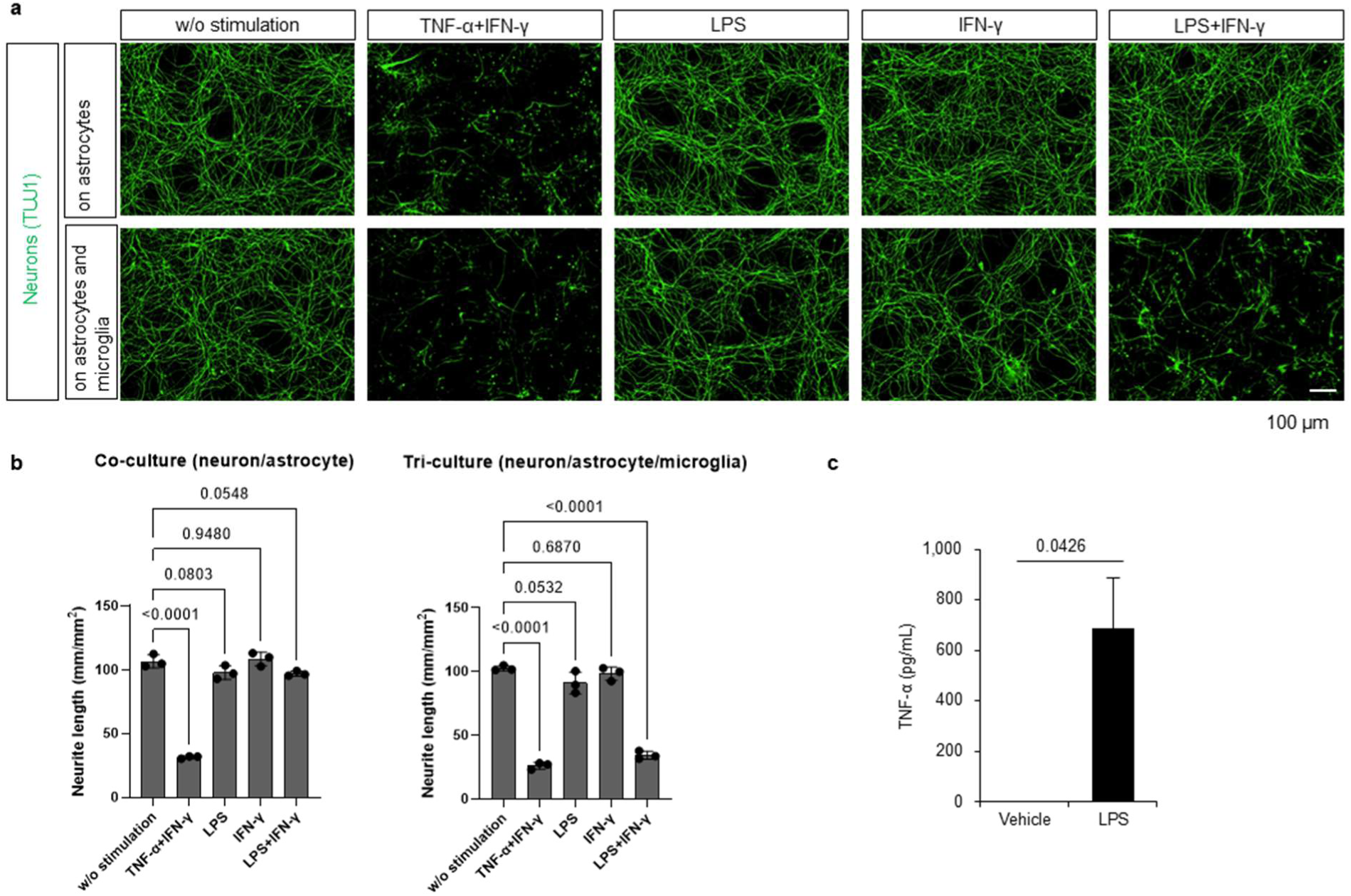
| LPS induces neurotoxicity in tri-culture in the presence of IFN-γ. **a,** Neurons on astrocyte or astrocyte/microglia cultures treated with TNF-α/IFN-γ, LPS alone, IFN-γ alone, or LPS/IFN-γ. Scale bar: 100 µm **b,** Quantification of TUJ1 positive neurite length of neurons on astrocytes (co-culture) or on astrocyte and microglia (tri-culture). Bar data represent mean ± s.d.; each dot represents an individual technical replicate (*n* = 3 wells). The reproducibility of the data was confirmed in an independent experiment. Statistical analysis was performed using one-way ANOVA followed by Dunnett’s multiple comparisons test, **c,** Levels of TNF-α released in the medium of tri-culture heated with LPS for 24 hours. The bar represents mean of four experiments with s.e.m. (*n* = 4 biological replicates). *P* values were computed using a two-sided *t*-test.

### Aged astrocytes induced stronger neurotoxicity in response to TNF-α and IFN-γ compared to young astrocytes

During our study, we noticed that some astrocytes induced strong neurotoxicity, depending on cell preparation lots, while others did not, even in the presence of both TNF-α and IFN-γ. Using commercially available astrocyte progenitor cells, we found that stronger neurotoxicity was induced when the progenitor cells were maintained for an extended period (approximately 3 months) before differentiating into astrocytes. We hypothesized that longer culture periods induced cellular senescence features and examined senescence markers, including γH2A.X, senescence-associated beta-galactosidase (SA-β-gal), and *CDKN2A* (p16) gene expression. As hypothesized, an increase in senescence marker expression was observed in astrocytes with a longer culture period (aged astrocytes), compared to astrocytes that were differentiated from astrocyte progenitor cells within 1 month (young astrocytes) (Fig. 4a, b). Long culture periods as differentiated astrocytes, which do not proliferate, did not increase *CDKN2A* expression (Extended Data Fig. 3), suggesting that these features might be caused by cellular senescence induced by cell proliferation stress in the astrocyte progenitor cell stage. Aged astrocytes with higher expression levels of senescence markers exhibited stronger neurotoxicity in response to TNF-α and IFN-γ compared to young astrocytes (Fig. 4c, d). Although the molecular mechanisms underlying the difference in TNF-α and IFN-γ response between young and aged astrocytes remain unclear, we observed that microglia cultured with aged astrocytes showed more rounded and less branched morphology compared to microglia cultured with young astrocytes (Fig. 4e, f). A reduced number of branches and a round shape are well-known characteristics of microglia in inflammatory reactive states ^11^. These results suggest that astrocytes undergo age-related changes, creating an inflammation-prone environment and becoming more sensitive to cytokines that induce a neurotoxic state.

**Fig. 4.**
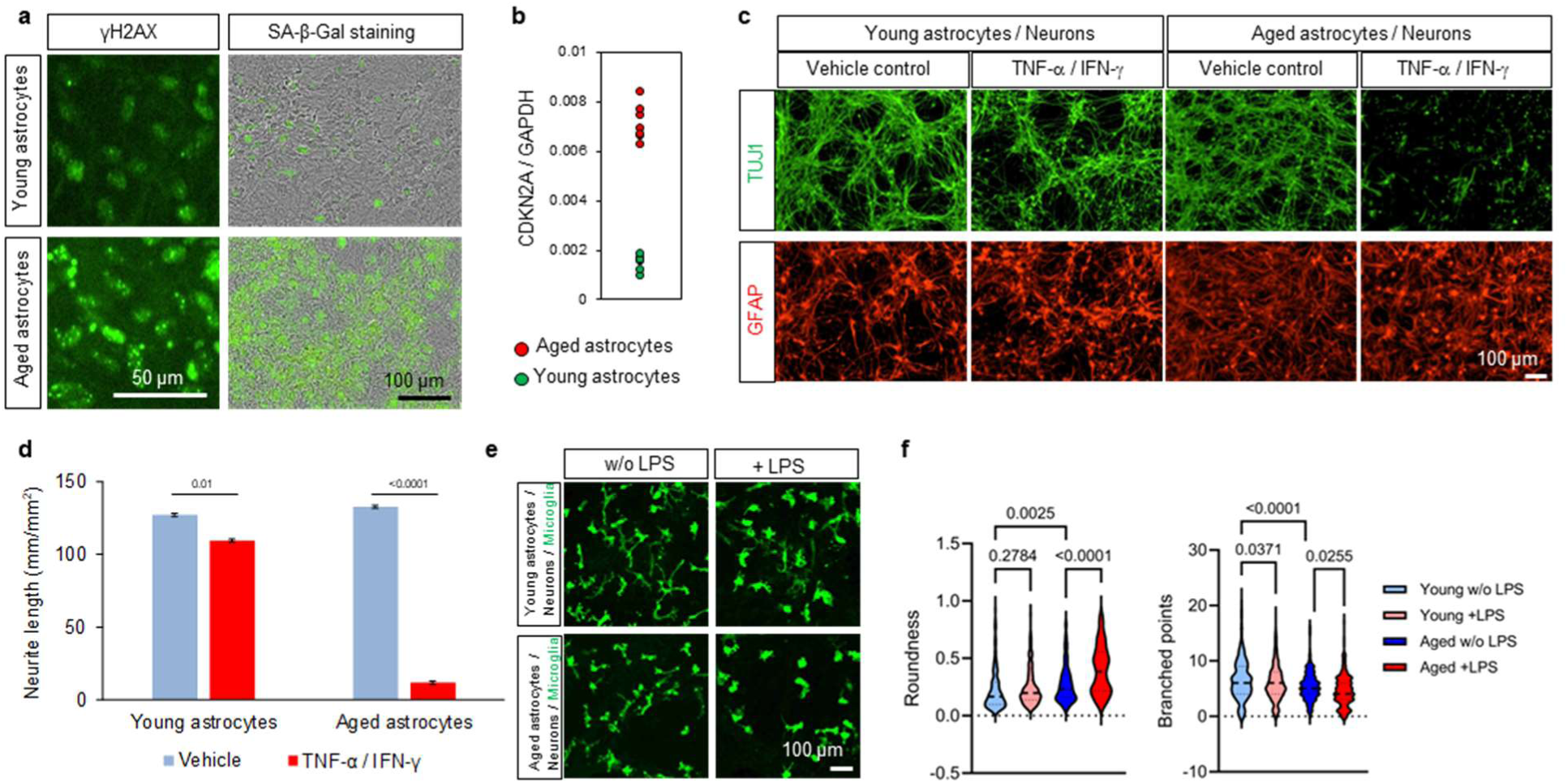
| Aged astrocytes induce strong neurotoxicity in response to TNF-α and IFN-γ. **a,** γH2AX and SA-β-Gal staining of human iPSC-derived astrocytes differentiated from astrocyte progenitor cells cultured for 1 month (young) or 3-4 months (aged), respectively, **b,** Relative levels of CDKN2A to GAPDH mRNA quantified by digital PCR. **c,** Neuronal (FUJI) culture on young or aged astrocytes (GFAP) treated with TNF-α/IFN- γ. **d,** Quantification of TUJ1+ neurite length in co-cultures **(c).** The bar indicates mean ± s.d. (*n* = 3 wells). *P* values were computed using a two-sided *t*-test. **e,** IBA1 staining to visualize morphology of microglia cultured on young or aged astrocytes in the presence or absence of LPS. **f,** Roundness and number of branched points of microglia. Violin plots show median and quartiles of cells (*n* = 406 cells for young astrocytes w/o LPS, 347 cells for young astrocytes with LPS, 261 cells for aged astrocytes w/o LPS, 220 cells for aged astrocytes with LPS). Statistical analysis was conducted using one-way ANOVA followed by Tukey’s multiple comparisons test.

### Astrocyte activation becomes more prominent following microglia activation in the aging human brain

To investigate the relationship between aging and the activation of microglia and astrocytes in the human brain, we analyzed publicly available data (GSE11882) from a study by Berchtold et al. ^12^ that examined gene expression changes during normal brain aging. The dataset includes gene expression profiles from the hippocampus of healthy donors ranging in age from 20 to 99 years. Using a method similar to that described previously ^13^, which groups genes based on temporal expression patterns, combined with dynamic Bayesian network analysis to model interactions between gene groups, we observed the activation of microglia-related genes became apparent around the ages of 30-40, followed by astrocyte activation around the ages of 50-60 (Fig. 5). In the gene groups where microglia and astrocyte genes are activated, activation of immune system-related genes was also recognized. These results suggest that microglia activate astrocytes and induce neuroinflammation. Furthermore, the time lag between microglia activation and astrocyte activation in the human brain may reflect the period during which astrocytes undergo aging.

**Fig. 5.**
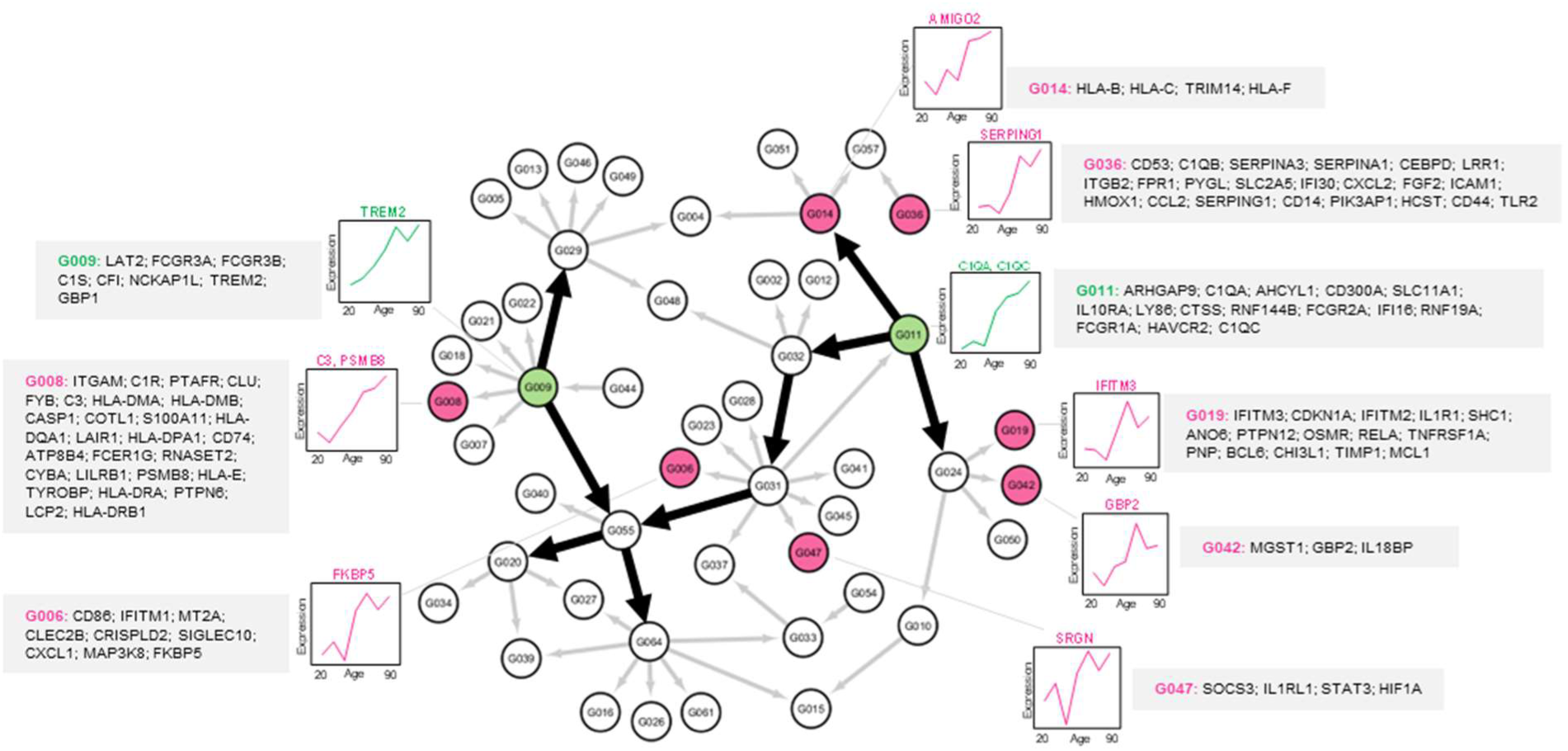
| Gene expression network analysis of aging brain reveals astrocyte activation following microglia activation. A gene expression network was constructed based on time-dependent pattern changes. Nodes containing genes related to microglia and astrocyte activation are colored green and pink, respectively. Within the nodes, genes associated with the immune system are listed.

## Discussion

Using human iPSC-derived astrocytes, we demonstrated that stronger neuronal toxicity was induced by TNF-α and IFN-γ compared to the combination of IL-1α, TNF-α, and C1q. We found that astrocytes expressing senescent markers responded more sensitively to TNF-α and IFN-γ, leading to neurotoxicity, whereas the astrocytes with low senescent marker expression showed a less prominent response. These results suggest the possibility that aging, in addition to inflammatory stimuli, is required to induce neuronal damage and may explain why neurodegenerative diseases are associated with aging.

Increased cellular senescence marker p16-positive astrocytes have been reported in the normal aged brain and in the brains of AD patients ^14^. While the molecular mechanisms by which aged astrocytes become more sensitive to inflammatory stimuli and/or induce neurotoxicity more strongly remains to be elucidated, it is known that astrocytes are remarkably impacted during aging, including the upregulation of genes associated with immune response and reactivity ^15^. It is also known that increased reactive astrocytes are recognized not only in disease brains but also normal aging brains ^16^. Our data, which indicate a likely higher activation of microglia on aged astrocytes compared to young astrocytes, may suggest that aged astrocytes create an environment prone to inflammation and receive additional signals further from activated microglia. It is also consistent with the gene network analysis showing increased expression of the genes related to inflammatory responses with aging.

Induction of C3 gene expression has been recently reported in human iPSC-derived astrocytes by the combination of IL-1α, TNF-α, and C1q ^17^ as well as in rodent astrocytes ^1^. Induction of neurotoxicity by these three factors has also been reported in human iPSC-derived astrocytes based on reduced neurite lengths, but it did not result in extensive loss of neurons ^17^. We found that the combination of TNF-α and IFN-γ induced higher expression of reactive astrocyte markers, including C3, and stronger neurotoxicity compared to the three factors IL-1α, TNF-α, and C1q in human astrocytes. Conversely, in rat primary astrocytes, the three factors IL-1α, TNF-α, and C1q induced stronger expression of reactive astrocyte markers than the combination of TNF-α and IFN-γ. These results suggest species difference in cytokine response between rodents and human in the induction of neurotoxic reactive astrocytes.

The release of TNF-α was induced in LPS-treated tri-cultures consisting of microglia, astrocytes, and neurons. In our previous study, we showed that microglia responded first to LPS and induced TNF-α, which in turn activated astrocytes and induced further TNF-α production in astrocytes ^18^. In contrast, IFN-γ induction was not detected in tri-cultures stimulated with LPS. It has been reported that T cells in aged brains express IFN-γ and they are clonally different from T cells in the blood ^19^. We speculate that IFN-γ may be supplied by T cells in the aged brain, which could contribute to the manifestation of neurodegenerative diseases with aging, in addition to astrocyte cellular aging.

The molecular mechanism of neurotoxicity remains to be elucidated. It may be due to active neuron-killing activity, loss of supportive function, or both. It has been reported that long-chain saturated fatty acids are upregulated in rodent reactive astrocytes and they mediate neurotoxicity ^20^. Lipidome analysis of human aged neurotoxic astrocytes and other future studies are expected to further enhance our understandings of mechanisms of neurotoxicity and neurodegenerative disease pathogenesis. Neurotoxicity was observed towards glutamatergic, motor, and dopaminergic neurons, suggesting that astrocyte neurotoxicity may be a common mechanism underlying neurodegenerative diseases including AD, ALS, and PD. Targeting astrocytes is expected to provide novel therapeutic approaches for neurodegenerative diseases.

To confirm the potential use of neurotoxic astrocytes in drug development, we conducted a screening for compounds that inhibit the induction of neurotoxicity and identified JAK inhibitors as the most frequent hits. JAK inhibitors have been identified as candidates for drug reposing in Alzheimer’s disease through a machine learning framework to study and predict the response of FDA-approved drugs ^21^. Baricitinib, a JAK inhibitor, is currently in phase 2 clinical trials ^22^. JAK inhibitors are also expected to be potential therapeutics for ALS ^23^. We expect that our study can contribute to a better understanding of neurotoxic astrocytes and neurodegenerative diseases, and the development of therapeutics.

## Methods

### Generation of human iPSC-derived astrocytes

For compound screening based on C3 expression, iCell Astrocytes (Fujifilm Cellular Dynamics cat. no. C1037) were used as human iPSC-derived astrocytes. For all other experiments, astrocytes were prepared using a serum-free method as described below. Human iPSC-derived astrocytes were generated from commercially available astrocyte progenitor cells (XCell Science cat. no. XCS-AP-001-1V, Axol Bioscience cat. no. AX0083, Applied Stem Cell cat. no. ASE-9322P). Astrocyte progenitor cells were seeded in poly-D-lysine (PDL)-pre-coated flask (Greiner Bio One) coated with iMatrox-511 silk (Matrixome), and maintained in the astrocyte progenitor medium containing DMEM/F12 (Thermo Fisher Scientific) supplemented with 1% N-2 Max Media Supplement (R&D Systems), 1% GlutaMAX Supplement (Thermo Fisher Scientific), 1% MEM Non-Essential Amino Acids Solution (Thermo Fisher Scientific), Penicillin-Streptomycin Solution (Fujifilm Wako), 5 IU/μL heparin (Fujifilm Wako), 10 ng/mL human basic fibroblast growth factor (bFGF, Thermo Fisher Scientific), 10 ng/mL human epidermal growth factor (EGF, Thermo Fisher Scientific). Cells were incubated at 37 °C and 5% CO_2_, and the culture medium was changed 3 times per week.

For astrocytes differentiation, the medium was changed 1 day after seeding of astrocyte progenitor cells to the astrocyte differentiation medium containing DMEM/F12 supplemented with 1% N-2 MAX Media Supplement, 1% GlutaMAX Supplement, 1% MEM Non-Essential Amino Acids Solution, 1% Penicillin-Streptomycin Solution, 0.5 U/mL heparin, 10 ng/mL Ciliary Neurotrophic Factor (CNTF, Thermo Fisher Scientific), and 10 ng/mL Bone morphogenetic protein 4 (BMP4, Thermo Fisher Scientific). Cells were incubated at 37 °C and 5% CO_2_, and astrocytes were used for experiments 5 days after initiating differentiation.

### Human iPSC-derived microglia and neurons

iCell Microglia (Fujifilm Cellular Dynamics cat. no. C1110) was used as human iPSC-dirved microglia. iPSC-derived neurons were purchased from Fujifilm Cellular Dynamics; iCell GlutaNeurons (cat. no. C1060, C1033), iCell GABAneurons (cat. no. C1008), iCell DopaNeurons (cat. No. C1087, C1028), and iCell Motor Neurons (cat No. C1048, C1050). NGN-2 induced cortical excitatory neurons, FF-1 and FF-2 iNeurons, were prepared from iPSCs as previously reported ^24^.

### Rat primary astrocytes

Fetal bovine serum (Thermo Fisher Scientific, cat. no. 10270-106) was heat-inactivated by incubation at 56°C for 30 minutes. Primary rat astrocytes (Thermo Fisher Scientific, Cat. No. N7745100) were seeded and maintained in the rat astrocyte medium containing DMEM, high glucose, no glutamine (Thermo Fisher Scientific) supplemented with 10% (v/v) heat-inactivated FBS. Cells were incubated at 37 °C and 5% CO_2_ for one week. The cells were passaged, and after a subsequent week of culture, cryopreserved stocks were prepared using Stem Cell Banker (Zenogen Pharma). These cryopreserved frozen stocks were used for subsequent assays using the serum-free rat astrocyte medium containing 50% Neurobasal medium (Thermo Fisher Scientific), DMEM/F12, 1% N-2 Max Media Supplement, 1% GlutaMAX Supplement, 1% Sodium Pyruvate (Thermo Fisher Scientific), Penicillin-Streptomycin Solution, 5 μg/mL *N*-acetyl cysteine (Sigma Aldrich) and 5 ng/mL HBEGF (Thermo Fisher Scientific).

### Immunostaining

Cells were fixed by adding formalin solution (Fujifilm Wako) and incubated for 30 minutes at room temperature. After washing with PBS (−), cells were permeabilized for 1 hour with PBS (−) containing 1% BSA (Sigma Aldrich) and 0.2% Triton X-100 (BioVision) (blocking solution). Cells were then incubated overnight at 4°C with primary antibody solution (anti-GFAP, rabbit, 1:5000, Merck Millipore; anti-TUJ1, mouse, 1:7000, Novus Biologicals; Anti-phospho-Histone H2A.X (Ser139) (γH2AX), mouse, 1:250, Merck Millipore; anti-IBA1, rabbit, 1:800, Fujifilm Wako). Following PBS (−) washes, cells were incubated for 1 hour at room temperature with secondary antibody solution (anti-mouse Alexa Fluor 555, goat, 1:1000, Thermo Fisher Scientific; anti-mouse Alexa Fluor 488, goat, 1:400-1000, Thermo Fisher Scientific; anti-mouse Alexa Fluor 594, rabbit, 1:1000, Thermo Fisher Scientific; anti-rabbit Alexa Fluor 488, goat, 1:1000, Thermo Fisher Scientific). After PBS (−) washes, images were captured using the IncuCyte® S3 Live-Cell Analysis System (Sartorius).

### RNA extraction and qRT-PCR

RNA extraction was performed using RNeasy Mini Kit (Qiagen), FastLane Cell SYBR Green Kit (Qiagen), or QIAseq UPX Cell Lysis Kit (Qiagen) according to the manufacturer’s instructions. qRT-PCR was conducted using FastLane Cell SYBR Green Kit or QuantiTect SYBR Green RT-PCR Kit (Qiagen), following the protocols provided by the manufacturer. The following primers were used: GAPDH (human), sense: 5’-gtcagtggtggacctgacct-3’, antisense: 5’-tgctgtagccaaattcgttg-3’; GAPDH (rat), sense: 5’-agaacatcatccctgcatcc-3’, antisense: 5’-gtcctcagtgtagcccagga-3’; C3 (human), sense: 5’-cacatcctatgccctcttgg-3’, antisense: 5’-ggcgtccttttggtattgag-3’; C3 (rat), sense: 5’-caactgtgtttcgcctgcta-3’, antisense: 5’-tctcaggggctggttttatg-3’; GBP2 (human), sense: 5’-tttcaccctggaactggaag-3’, antisense: 5’-gacgaagcacttcctcttgg-3’; GBP2 (rat), sense: 5’-gaacaaattggctgggaaga-3’, antisense: 5’-tgtagacgaaggtgctgctg-3’; PSMB8 (human), sense: 5’-gctattctggaggcgttgtc-3’, antisense: 5’-aggcctcttcttctccttgg-3’; PSMB8 (rat), sense: 5’-tctcgggacagatgttctcc-3’, antisense: 5’-cactttcacccaaccgtctt-3’; FBLN5 (human), sense: 5’-ccctactcgaccccctactc-3’, antisense: 5’-gttgcacactcgtccacatc-3’; FBLN5 (rat), sense: 5’-ggccctactccaatccctac-3’, antisense: 5’-gtaccctccttccgtgttga-3’; SERPING1 (human), sense: 5’-gggggatgctttggtagatt-3’, antisense: 5’-caggtgaagtccttggggta-3’; SERPING1 (rat), sense: 5’-cgaagaaggctgagaccaac-3’, antisense: 5’-gtggacacaggcaaaatcct-3’; CXCL10 (human), sense: 5’-aggaacctccagtctcagca-3’, antisense: 5’-caacacgtggacaaaattgg-3’; CXCL10 (rat), sense: 5’-gcttattgaaagcggtgagc-3’, antisense: 5’-ggtcaggagaaacagggaca-3’; Cycrophilin B (human), sense: 5’-gcacaggaggaaagagcatc-3’, antisense: 5’-agccaggctgtcttgactgt-3’; JAK1 (human), sense: 5’-agcgatgtccttaccacacc-3’, antisense: 5’-cctcaacacactcaggagca-3’; P2RY4 (human), sense: 5’-ctggactgttggtttgatgagga-3’, antisense: 5’-cagcgacagcacatacaaggt-3’). qRT-PCR was conducted using the CFX384 Touch Real-Time PCR Detection System (Bio-Rad). The data were analyzed using the delta-delta Ct method.

### Neurotoxicity assay

iPSC-derived astrocytes were prepared at a density of 50,000-60,000 cells per well (96 well plate), or 10,000-12,000 cells per well (384 well plate). For astrocytes and microglia co-culture experiments, iCell Microglia were seeded at 30,000 cells per well (96 well plate), or 6,000 cells per well (384 well plate). Subsequently, cultures were stimulated for 2 days with 7.5 ng/mL TNF-α (Cell Signaling Technology) and 5-7.5 ng/mL IFN-γ (R&D Systems), or 100 ng/mL LPS (Fujifilm Wako) and 5 ng/mL IFN-γ. Alternatively, cells were treated with 3 ng/mL IL-1α, (Sigma Aldrich), 30 ng/mL TNF-α, and 400 ng/mL C1q (Abcam) for 2 days in a condition reported by Liddelow et al. (Liddelow et al., 2017). For siRNA knockdown experiments, cells were treated with Accell siRNA (Horizon Discovery) at a final concentration of 3 μM for 7 days. This treatment was repeated after 1 week, followed by stimulation with 7.5 ng/mL TNF-α and 5 ng/mL IFN-γ for 2 days. Following stimulation, the culture medium was removed, and neurons were seeded at a density of 30,000 cells per well (96 well plate), or 6,000 cells per well (384 well plate). After 3 days, cells were fixed and immunostained for TUJ1. Subsequently, neurite length was quantified using the IncuCyte® S3 Live-Cell Analysis System with NeuroTrack Software module (Sartorius).

### Compound screening targeting C3 expression

Compound solutions (10 mM) for screening were purchased the FDA-approved Drug Library (Selleck Chemicals), Inhibitors, Agonists, Screening Libraries (MedChemExpress), and StemSelect Small Molecule Regulators 384-Well Library I (Merck Millipore). iCell astrocyte were seeded in 384-well plates coated with Matrigel (Corning) at a density of 6,000 cells per well. One day after seeding, cells were stimulated for 48 hours with compounds at a final concentration of 10 μM (0.1% v/v), along with 7.5 ng/mL TNF-α and 5 ng/mL IFN-γ. RNA was subsequently extracted using the FastLane Cell Multiplex Kit (Qiagen), and qRT-PCR was performed using either the FastLane Cell Multiplex Kit or the QuantiTect Multiplex RT-PCR NR Kit (Qiagen). The following primers (Takara Bio) were used: GAPDH, sense: 5’-gaaccatgagaagtatgacaacagc-3’, antisense: 5’ tgggtggcagtgatggca-3’, probe: 5’-HEX-catcagcaatgcctcctgcaccaccaa-BHQ1-3’; C3, sense: 5’-tggaatctctacgaagctcatgaa-3’, antisense: 5’-cctgcattactgtgacctcgaa-3’, probe: 5’-FAM-atgtcggacaagaaagggatctgtgtgg-BHQ1-3’. PCR was conducted using the CFX384 Touch Real-Time PCR Detection System. C3 mRNA expression levels were quantified using the delta-delta Ct method using GAPDH as the endogenous control.

### V-PLEX

iPSC-derived astrocytes were prepared at a density of 60,000 cells per well in 96 well plates, Subsequently, iCell Microglia and induced neurons derived from iPSCs were seeded on the astrocytes at a density of 30,000 cells per well each. After approximately two weeks of co-culture, the cells were stimulated with 100 ng/mL LPS. After 24 hour, culture supernatants were collected and centrifuged at 2,000×g, 5 minutes at room temperature. Cytokine quantification was conducted using the V-PLEX Human Proinflammatory Panel II (4-Plex) kit (Meso Scale Discovery, MSD), following the manufacturer’s protocol as described in our previous report ^18^. Samples were diluted 10-fold with Diluent 2 prior to measurement. Signal detection was performed using the Meso QuickPlex SQ 120 MM instrument (MSD).

### SA-β-Gal staining

SA-β-Gal staining was performed using the Cellular Senescence Detection Kit -SPiDER-βGal (Dojindo) according to the manufacturer’s protocol. Briefly, Mcllavine buffer (pH 6.0) was prepared by mixing 3.7 mL of 0.1 mol/L citric acid solution with 6.3 mL of 0.2 mol/L disodium hydrogen phosphate solution. Astrocytes fixed with formalin solution were washed three times with Hanks’Balanced Salt Solution (HBSS) (Fujifilm Wako). The SPiDER-βGal DMSO stock solution was diluted 2,000-fold in Macllvaine buffer and applied to the cells. After incubation at 37°C for 30 minutes, the cells were washed tree times with HBSS and then images were acquired using IncuCyte® S3 Live-Cell Analysis System.

### Droplet digital PCR

RNA extraction was performed using the RNeasy Mini Kit according to the manufacturer’s protocol. Reverse transcription was carried out using the PrimeScript RT reagent Kit with gDNA Eraser (Takara Bio) following the manufacturer’s protocol. Droplet digital PCR was conducted using the ddPCR EvaGreen Supermix (Bio-Rad) and QX200 AutoDG Droplet Digital PCR system (Bio-Rad) according to the manufacturer’s protocol. Briefly, cDNA prepared by reverse transcription was diluted 25-fold for *CDKN2A* detection and 500-fold for *GAPDH* detection. Diluted samples were added the reaction mixture contained 1× ddPCR EvaGreenSupermix and 136 nM of each forward and reverse primer. The following primers were used: GAPDH (Human Housekeeping Gene Primer Set, Takara Bio); CDKN2A (Takara Bio), sense: 5’-gcacattcatgtgggcatttc-3’, antisense: 5’-agctttggttctgccatttgcta-3’. Then, Droplets were generated using the Automated Droplet Generator (Bio-Rad), and PCR amplification was performed using the C1000 Touch Thermal Cycler (Bio-Rad). Subsequently, droplets were analyzed using the QX200™ Droplet Reader (Bio Rad) for mRNA quantification.

### EVs extraction and ELISA

Culture supernatants were collected from astrocytes treated with 7.5 ng/mL TNF-α and 5 ng/mL IFN-γ, or vehicle-treated astrocytes. Exosomes were isolated from these supernatants using the MagCapture Exosome Isolation Kit PS (Fujifilm Wako) according to the manufacturer’s protocol. The quantity of EVs was relatively quantified using the CD9/CD63 Exosome ELISA Kit (Cosmo Bio). Briefly, a calibration curve was generated using CD9/CD63 fusion protein as the standard, and the relative quantity of EVs in the samples was determined based on this standard curve. The amount of complement C3 in the EV samples was quantified using the Complement C3 Human ELISA Kit (Abcam). To evaluate the C3 content per EV, the total C3 concentration was normalized to the relative EV quantity in each sample.

### Imaging analysis of microglial morphology

Image analysis was performed on two–channel micrographs comprising a nuclear–staining channel and an IBA1 channel that selectively labels microglia. All computations were executed in Python 3.7.9; semantic segmentation networks were implemented in Keras (TensorFlow backend) and the thinning routine relied on the skeletonize function of scikit-image. Nuclear objects were first identified with a UNet-based model that assigns every pixel to one of three classes—nucleus, nuclear center or background—thereby providing both binary nuclear masks and discrete nuclear-center coordinates in a single forward pass. The microglial channel was independently processed with a second UNet that classifies pixels as soma, microglial processes or background. Both networks had been trained in advance on image sets acquired independently of the material analyzed here.

The preliminary soma mask was cross-checked against the nuclear mask. Any soma label devoid of an underlying nuclear signal was re-classified as microglial process, effectively eliminating false-positive somas. The validated soma and process regions were then skeletonized; branch points and termini of the resulting one-pixel-wide skeleton were regarded as graph nodes and the intervening skeletal segments as edges, yielding an undirected spatial graph for each microglial cell.

Nodes and edges located inside the soma were subsequently removed, and each soma was re-inserted into the graph as a single node positioned at its nuclear center. If a graph contained more than one nuclear-center node, neighboring microglia were considered to have been merged erroneously; in such cases, the graph was partitioned by severing the shortest path between individual nuclear centers until each sub-graph contained exactly one nucleus. For every cell-specific graph thus obtained, quantitative descriptors were extracted: the geometry of microglial processes (number of branch points, and related metrics).

### Gene expression analysis of human brain aging

Publicly available gene expression data related to human brain aging were analyzed with a modified version of the method described in a previous report ^13^. Briefly, a gene expression dataset (GSE11882; ^12^) derived from the hippocampi of cognitively intact individuals aged 20 to 99 years was analyzed to investigate human brain aging. Samples were divided into decade-based age groups, but the 50s group was excluded due to insufficient sample size. The data were normalized using the MAS5 algorithm, from which 14,046 unique genes with the highest JetSet scores ^25^ were selected.

Using a spline-based algorithm ^26^, we identified 712 genes that were differentially expressed across the seven age groups (q-value < 0.01). These genes were subsequently clustered into 64 groups based on their temporal expression patterns using hierarchical clustering with Pearson’s correlation as the distance metric. Mean expression values were calculated for each group, and interactions among groups were modeled with a dynamic Bayesian network ^27^, resulting in a network of 48 connected nodes after filtering edges with bootstrap probabilities < 0.5. The analysis was performed on the Shirokane supercomputer and visualized with Cytoscape ^28^.

#### Data source

A gene expression dataset related to human brain aging was obtained from the NCBI Gene Expression Omnibus (GSE11882; ^12^). This dataset was generated using the Affymetrix Human Genome U133 Plus 2.0 Array and includes samples derived from the post-mortem hippocampi of 43 cognitively intact individuals, ranging in age from 20 to 99 years. The samples were categorized into eight decade-based age groups, spanning individuals in their 20s through their 90s. However, due to the availability of only a single sample for the 50s age group, it was excluded from subsequent analyses since meaningful statistical evaluations could not be performed. As a result, seven age groups, each consisting of 4 to 9 samples, were retained for downstream analyses.

#### Preprocessing

The expression data were normalized using the MAS5 algorithm implemented in the Affymetrix Expression Console software. A total of 26,400 probe sets were identified based on two criteria: (i) they exhibited suprathreshold signals with a “present” call across all samples within at least one of the seven age groups, and (ii) they were assigned a valid gene symbol. For genes represented by multiple probe sets, the probe set with the highest JetSet score ^25^ was selected as the representative, while the others were excluded. This process yielded a dataset of 14,046 unique genes. Expression values were subsequently log2-transformed for further analyses.

#### Differential expression filtering

Genes exhibiting differential expression across the seven age groups were identified using a spline-based algorithm implemented in the EDGE software ^26^. A total of 712 differentially expressed genes were identified by applying a q-value threshold of 0.01.

#### Grouping genes by temporal expression pattern

Next, the 712 differentially expressed genes were classified into 64 groups based on their temporal expression patterns. This grouping was performed using hierarchical clustering across the seven age groups, with the distance metric based on Pearson’s correlation coefficient. For each group, an average expression value was calculated by taking the mean of the expression values of its member genes.

#### Network modeling for groups of genes

The interactions among the 64 gene groups were modeled using a dynamic Bayesian network approach. The SiGN-BN software ^27,29^ was applied to the three-dimensional expression matrix [64 groups × 7 time points (age groups) × 4 to 9 samples], resulting in a network graph in which nodes (vertices) represent gene groups and edges represent their inferred regulatory dependencies. The analysis was performed with 10,000 pseudo-bootstrap replications, and edges with a bootstrap probability of less than 0.5 were excluded. The final network comprised 48 nodes connected by at least one edge, while unconnected nodes were removed. The SiGN-BN computations were carried out on the Shirokane supercomputer at the University of Tokyo. The resulting network was visualized using Cytoscape 3.10.3 software ^28^.

#### Pathway enrichment analysis

The genes in each node were analyzed for enriched pathways using Reactome pathway database ^30^.

## Acknowledgements

The authors would like to thank Kazuhiko Nakata, Ayako Sakai, Naohiro Iwakura, and Mitsuho Taniguchi for their assistance in the experiments. The authors also would like to thank colleagues of Fujifilm corporation for helpful discussions and support.

## Author contributions

HKo: conceptualization, investigation, formal analysis, writing - original draft. KM: conceptualization, formal analysis. TW: formal analysis. KK: conceptualization. AN: conceptualization, writing - review and editing. ES: investigation, HKa: conceptualization. S E-Y: conceptualization, formal data analysis, writing - original draft, writing - review and editing, supervision, project administration.

---

**Extended Data Fig. 1:**
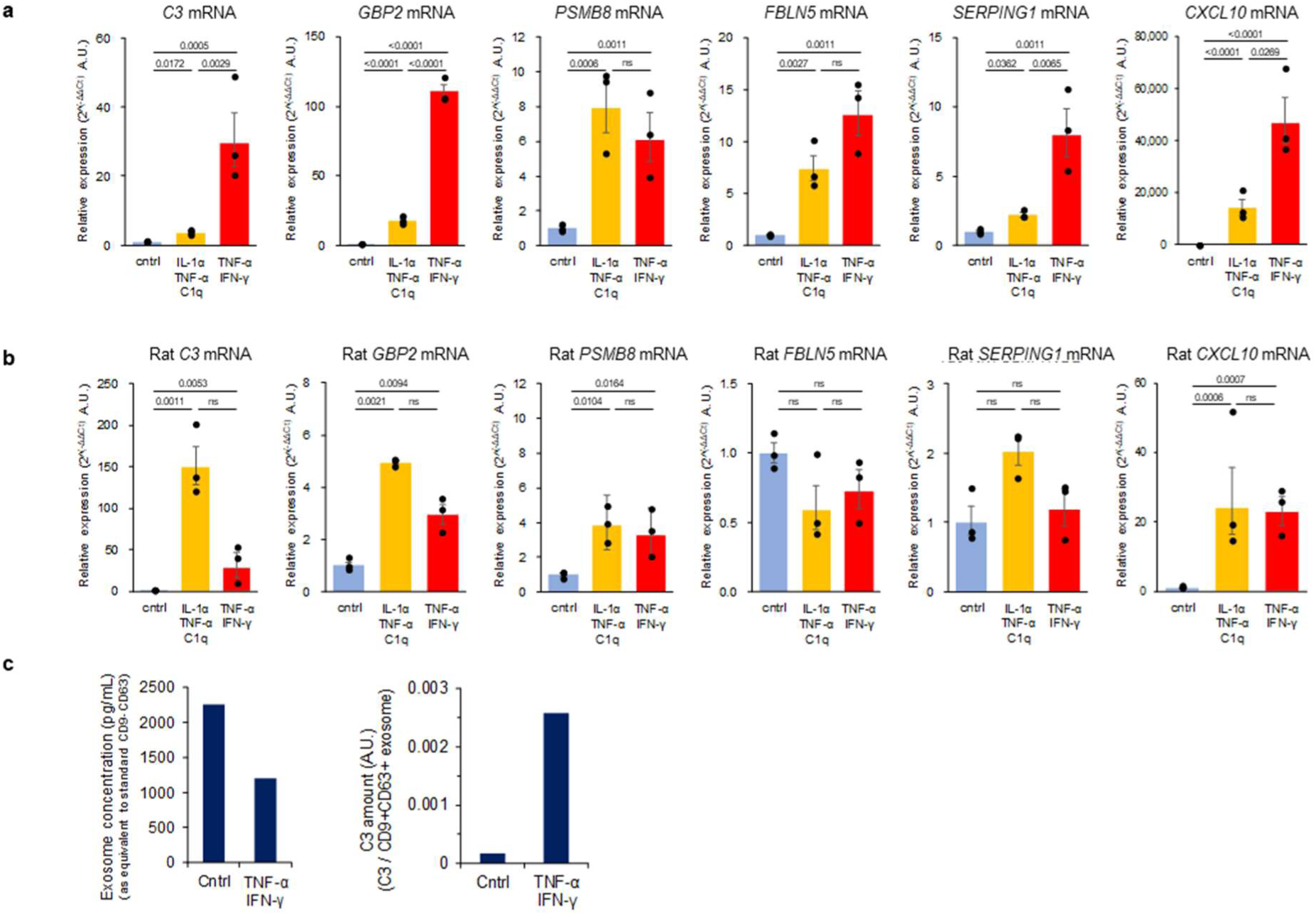
Astrocyte response to combination of TNF-α and IFN-γ. **a-b,** Relative gene expression levels of *C3, GBP2, PSMB8, FBLN5, SERPING1,* and *CXCL10,* using *GAPDH* as the internal control in human iPSC-derived astrocytes **(a)** and in rat primary astrocytes **(b)** cultured in serum-free rat astrocyte medium. Bar data represent mean ± s.e.m.; each dot represents an individual biological replicate (*n* = 3 independently prepared cells). Statistical analysis was performed using two-way ANOVA followed by Tukey’s multiple comparisons test; ns: 0.05 < *P.* **c,** Quantification of exosome concentration and C3 levels in exosomes isolated from the culture medium of human iPSC-derived astrocytes in the absence or presence of TNF-α and IFN-γ.

**Extended Data Fig. 2:**
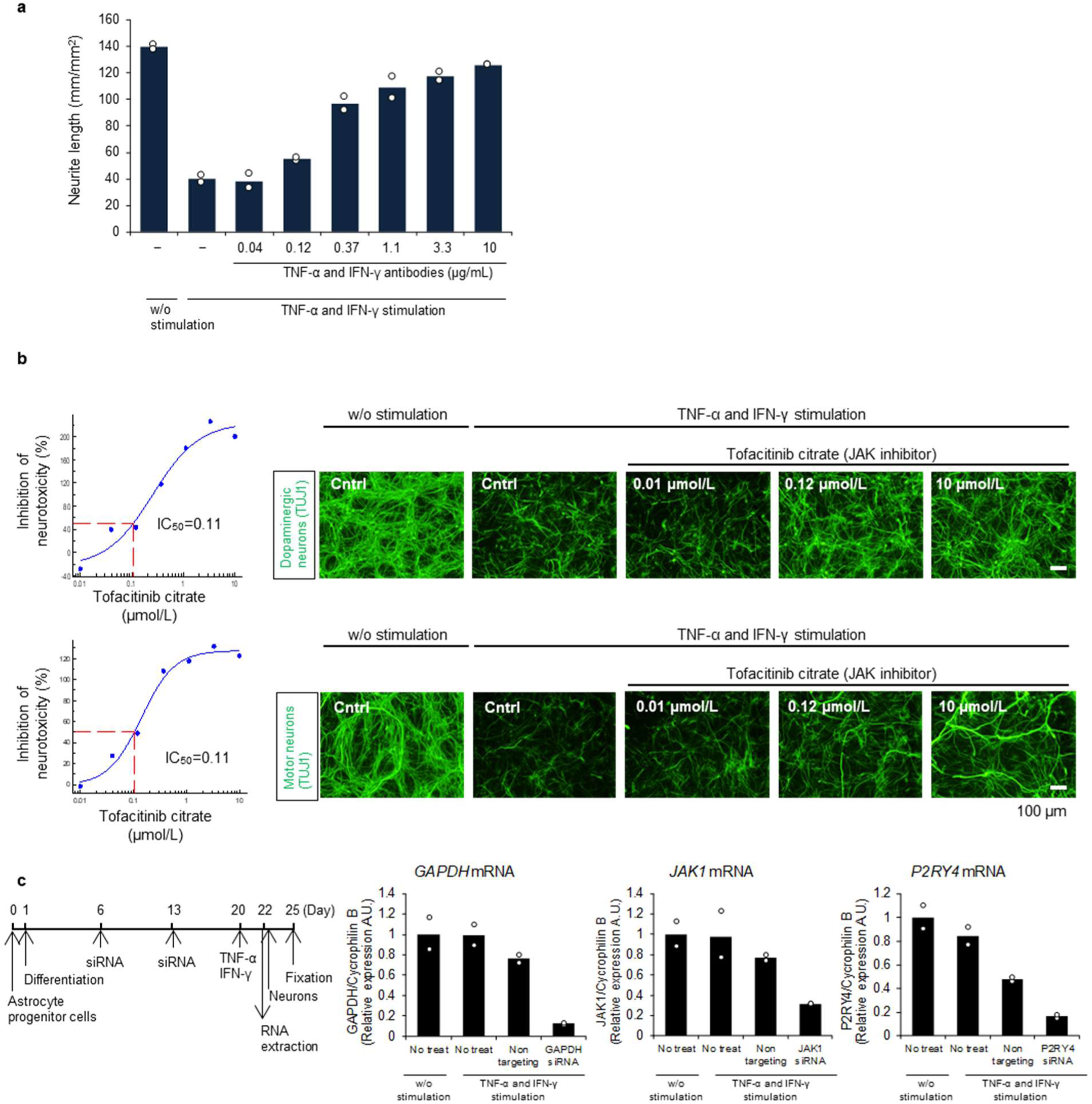
Neurotoxicity assay and compound treatment. **a,** Neurite length was analyzed by measuring TUJ1-positive neurites of neurons cultured on human iPSC- derived astrocytes. The astrocytes were treated with TNF-α and IFN-γ antibodies alongside TNF-α and IFN- γ. Bar data represent the average of 2 wells. Each dot represents an individual technical replicate (*n* = 2 wells), **b,** Neurotoxicity assay of astrocytes treated with Tofacitinib citrate using dopaminergic neurons and motor neurons. The dose-responsive curve is presented as the mean of *n* = 2 technical replicates, **c,** Schematic of gene knockdown using siRNA and confirmation of knockdown efficiency by qRT-PCR for *GAPDH. JAKI,* and *P2RY4*mRNA. Bar graphs represent the average of 2 wells, and each dot represents an individual technical replicate (*n* = 2 wells).

**Extended Data Fig. 3:**
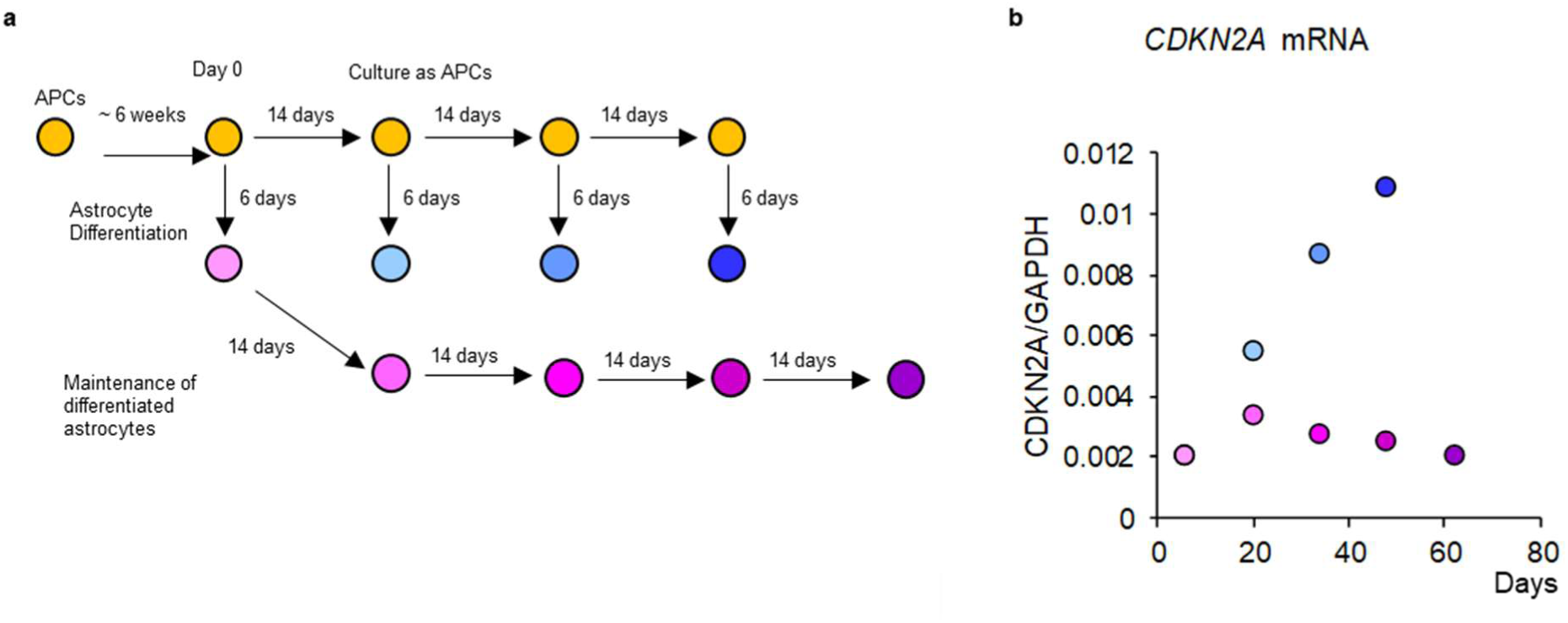
Astrocyte progenitor culture duration and induction of *CDKN2A*. **a,** Schematic of culture of astrocyte progenitor cells (APCs), differentiation into astrocytes, and maintenance as astrocytes, **b,** Relative levels of *CDKN2A* to *GAPDH* mRNA quantified by digital PCR in astrocytes, which were differentiated after culture duration as APCs (blue series of dots), or maintained as astrocytes (park series of dots).

